# *In Vivo* Simvastatin and Brain Radiation in a Model of HER2^+^ Inflammatory Breast Cancer Brain Metastasis

**DOI:** 10.1101/2024.05.25.595905

**Authors:** Swaminathan Kumar, Richard A. Larson, Shane Stecklein, Jay Reddy, Bisrat G. Debeb, Richard A. Amos, Stephanie M. Cologna, Wendy A. Woodward

## Abstract

**Purpose:** Inhibiting HMG-CoA reductase with simvastatin prevents breast cancer metastases in preclinical models and radiosensitizes monolayer and stem-like IBC cell lines *in vitro*. Given the extensive use of simvastatin worldwide and its expected penetration into the brain, we examined whether regulating cholesterol with simvastatin affected IBC3 HER2+ brain metastases.

**Methods and Materials:** Breast cancer cell lines KPL4 and MDA-IBC3 were examined in vitro for DNA repair after radiation with or without statin treatment. Brain metastasis endpoints were examined in the MDA-IBC3 brain metastasis model after ex vivo exposure to lipoproteins and after tail vein injections with and without whole-brain radiotherapy (WBR) and oral statin exposure.

**Results:** *Ex vivo* preculture of MDA-IBC3 cells with very low-density lipoprotein (vLDL) enhanced the growth of colonized lesions in the brain *in vivo* compared with control or high-density lipoprotein (HDL), and concurrent oral simvastatin/ WBR reduced the incidence of micrometastatic lesions evaluated 10 days after WBR. However, statin, with or without WBR, did not reduce the incidence, burden, or number of macrometastatic brain lesions evaluated 5 weeks after WBR.

**Conclusions:** Although a role for cholesterol biosynthesis is demonstrated in DNA repair and response to whole brain radiation in this model, durable in vivo efficacy of concurrent whole brain irradiation and oral statin was not demonstrated.

## INTRODUCTION

Inflammatory breast cancer (IBC) typically appears as a red and swollen breast with changes in the overlying skin, with or without a palpable mass (reviewed in(*1*)). It is the most lethal form of breast cancer; indeed, approximately 30% of patients with IBC have metastatic disease at the time of diagnosis. The cumulative incidence of brain metastases in IBC at 2 years is 19%(*2*). The risk increases further for breast cancer of specific receptor subtypes: patients with human epidermal growth factor receptor 2-enriched (HER2^+^) or triple-negative stage IV breast cancer are at a 25%-45% risk of developing brain metastasis(*1, 3, 4*).

Despite improvements in multimodal therapy, including new agents with activity against HER2^+^ brain metastases, progression-free survival rates at 1 year after the appearance of brain metastases are low(*5*). In this study we used a previously described HER2^+^ MDA-IBC3 mouse model of metastatic breast cancer, in which tail-vein injection of HER2^+^ IBC cells leads to a high rate of brain metastasis, to examine strategies for radiosensitizing developed lesions(*6*). We hypothesized that the combination of simvastatin and WBR may synergize to eradicate established IBC-3 brain metastases *in vivo because* patients with IBC taking statins (i.e., inhibitors of HMG-CoA reductase and cholesterol biosynthesis) have been reported to have better primary tumor control and better recurrence-free survival than patients who did not take statins(*7*), published evidence suggest that cholesterol modulation by lipoproteins and statins regulates radiosensitivity in IBC(*8–11*), and re-analysis of gene expression studies described from MDA-IBC3 sublines(*6*) demonstrate brain metastases express cholesterol regulation genes more strongly than do lung metastases.

## MATERIALS & METHODS

### Cell culture

MDA-IBC HER2^+^ green fluorescent protein (GFP) expressing cells were used for *in vivo* experiments(*6*). Cells were cultured in Ham’s F-12 medium supplemented with 10% fetal bovine serum (FBS), 1 µg/mL hydrocortisone, 5 µg/mL insulin, and 1% antibiotic-antimycotic, and maintained in a humidified, 5% CO_2_ environment at 37°C. Lipoprotein experiments used serum-free medium to avoid effects from the cholesterol and lipoprotein in FBS. IBC3 cells were cultured for 24 hours with very low-density lipoprotein (vLDL) or high-density lipoprotein (HDL, Sigma-Aldrich, St. Louis, MO) at 10 μg/mL.

### Traffic Light Reporter Assay

We used a Traffic Light Reporter (TLR) assay, described elsewhere (*12*), to assess simultaneous homologous recombination [HR] and non-homologous end joining [NHEJ] in IBC cells. Briefly, the cargo plasmids pCVL Traffic Light Reporter 1.1 (Sce Target) Ef1a BFP (cat #31481), pCVL SFFV EF1s HA.NLS.Sce(opt) (cat #31479), pCVL SFFV d14GFP Donor (cat #31475), and pCVL SFFV d14GFP EF1s HA.NLS.Sce(opt) (cat #31476) were obtained from Addgene. Purified plasmids were transfected into HEK293 cells along with two lentiviral packaging plasmids, pCMV-VSVG (Addgene #8454) and psPAX2 (Addgene #12260), by using Lipofectamine 2000 (Invitrogen). Viral supernatants were collected and concentrated with a Lenti-X concentrator (Takara) and stored at –80°C. To generate reporter cell lines, we infected two breast cancer cell lines, KPL4 (preferentially metastasizes to lymph nodes and lungs) and IBC3 (metastasizes preferentially to brain), with lentivirus produced with the pCVL Traffic Light Reporter 1.1 (Sce Target) Ef1a BFP cargo plasmid, and 72 later BFP^+^ cells were sorted on a FACSAria II cell sorter (BD Biosciences). Sorted KPL4^TLR-BFP^ and IBC3^TLR-BFP^ cells were expanded and cryopreserved. For the TLR assay, 2.5×10^5^ KPL4^TLR-BFP^ or IBC3^TLR-BFP^ cells were plated in each well of a 6-well tissue culture plate and incubated overnight to allow cells to attach. The following day, fresh medium supplemented with vehicle, the HMG-CoA reductase inhibitor simvastatin (10 µM in DMSO), (R)-mevalonic acid (an intermediate in cholesterol synthesis; the product of HMG-CoA reductase); 100 µM in ethanol), or a combination of simvastatin and (R)-mevalonic acid was added. At 24 hours, fresh medium supplemented with vehicle, simvastatin, (R)-mevalonic acid, or a combination of simvastatin and (R)-mevalonic acid with either no virus (control), the pCVL SFFV EF1s HA.NLS.Sce(opt) (“I-SceI”) virus, the pCVL SFFV d14GFP Donor (“GFP”) virus, or the pCVL SFFV d14GFP EF1s HA.NLS.Sce(opt) (“I-SceI/GFP”) virus was added and the cultures incubated for another 48 hours. Cells were then trypsinized and analyzed for GFP and mCherry expression on a Gallios flow cytometer (Beckman Coulter). Data were analyzed in FlowJo V10.

### G1-to-S Transition Assay

KPL4 cells (1×10^5^) and IBC3 cells (5×10^5^) were plated in 6-well plates in complete medium and allowed to attach overnight. The next day, the medium was aspirated, cells were washed in 1X phosphate-buffered saline (PBS), and then incubated in serum-free medium for 24 hours. The medium was replaced with serum-free medium supplemented with either vehicle or simvastatin for an additional 21 hours. The medium was then replaced with complete medium (10% FBS) supplemented with vehicle or simvastatin as well as 10 µM 5-ethinyl-2’-deoxyuridine (EdU) and 1 µg/mL colcemid. Cells were returned to the incubator for 24 hours and then processed for EdU incorporation by using a Click-iT EdU Alexa Fluor 647 flow cytometry kit (Invitrogen) and a LSR II flow cytometer (BD Biosciences). Data were analyzed in FlowJo V10.

### Tail-vein injection

Three-to five-week-old female immunocompromised SCID/Beige mice (Harlan, USA) were housed and used in accordance with the institutional guidelines of The University of Texas MD Anderson Cancer Center under protocols approved by the Institutional Animal Care and Use Committee. MDA-IBC3 cells labeled with GFP were cultured as described above until 60%-70% confluence, treated with 0.25% trypsin-ethylenediaminetetraacetic acid (EDTA), counted (5×10^5^ cells in 200 μL PBS per mouse), and intravenously injected into 6-week-old mice via the tail vein (30-gauge needle). Mice were monitored regularly and weighed weekly after tumor-cell injection. Mice were euthanized and subjected to necropsy at 8-9 weeks after injection of the tumor cells.

### Statin and whole-brain radiotherapy treatment

*In vivo* experimental schemas are presented in Supplementary Figure S1. At 3 or 4 weeks after cell injection, mice were irradiated with 10 Gy in one fraction or 9 Gy in three 3-Gy fractions according to the experimental plan. For radiation treatment planning and delivery, mice were anesthetized with isoflurane and placed in the imaging and treatment stage of an X-RAD 225Cx small-animal irradiator (PRECISION X-RAY, North Branford, CT, USA); cone-beam computed tomography images were obtained with a 2.0-mm aluminum filter at 40 kVp and 2.50 mA and used to manually set the isocenter for each mouse. All mice were irradiated according to the same treatment plan with opposed lateral beams, developed with PilotXRAD 1.10.4 software. Dose was prescribed to the isocenter. Isocenter placement was selected to exclude the aerodigestive tract. Irradiation was done with a 0.3-mm copper filter at 225 kVp and 13.0 mA, with a 15-mm-diameter field size, at a dose rate of approximately 3.2 Gy/min.

For experiments assessing the effect of simvastatin, mice began receiving water supplemented with simvastatin and refreshed weekly at a concentration of 0.06 mg/mL as described previously(*11*).

### Fluorescent microscopy

Metastatic status was determined by fluorescent microscopy (SMZ1500, Nikon Instruments, Melville, NY). (*6*) Endpoints assessed were incidence (a binary variable, that is, presence or absence) of metastases in each brain; numbers of metastases in the brain, and tumor burden per brain (global GFP assessment). Three researchers blinded to treatment group and not involved in imaging counted the number of brain metastatic lesions per mouse in GFP images of the top and bottom of the brain. For the incidence of brain metastatic lesions, the three researchers’ counts were plotted individually. For the number of brain metastatic lesions, the arithmetic mean of three researchers’ counts were plotted for each treatment group. If two metastatic lesions coalesced without clear separation, they were counted as one. For tumor burden, the threshold between actual GFP signal and background was set initially with ImageJ software. The same threshold intensity value was applied for all of the images automatically by the ImageJ macro program. Tumor burden was calculated by dividing the integrated (top and bottom) GFP pixel area by the integrated total brain area.

### Mass spectrometry-based tissue imaging

For sample preparation, brain tissue with GFP-positive metastatic lesions (IBC3-GFP cells) were sectioned into slabs with a scalpel, flash-frozen with dry ice, and then stored at –80°C. The frozen tissue was cut into 10-µm-thick sections with a cryotome, and 31 such serial sections were placed on 31 glass microscope slides. Every 5^th^ slide (slide no. 1, 6, 11, 16, 21, 26, and 31) was stained with hematoxylin and eosin (H&E) to verify the presence of brain metastatic lesions. Slides with at least 1 metastatic lesion were chosen for tissue mass spectrometry–based imaging. Those slides were kept frozen at –80°C until ready for matrix application. A Shimadzu IM Layer, an automated sublimation matrix applicator, was used to coat the tissue sections with 2, 5 dihydroxybenzoic acid (for positive mode) for 30 min. The coated slides were then rehydrated in a heated humidifying chamber for 3 minutes with 1 mL of 9:1 water: methanol solution.

The cryo-sectioned slides were imaged by a Waters Synapt G2-Si with imaging MALDI mass spectrometer. Tissues mounted on microscope slides were imaged with a 2.5-kHz NdYAG solid state laser rastered across the tissue sample, providing a chemical composition profile for each corresponding spatial coordinate. This mass spectral information was collated by High-Definition Imaging (HDI) software to produce a chemical image that can be correlated with the sample’s histological profile (from H&E staining of adjacent tissue slide). Before the coated slide was loaded into the mass spectrometer, the slide was scanned with an EPSON scanner to map the areas of interest into the HDI software. The laser power was set to 250 (arb units) with 300 laser shots per pixel data. The angle of incidence of the laser projected elliptical spots with major axis length of 60 µm onto the sample plane, and the sample raster step was set to 60 µm accordingly. Before data were acquired, the instrument was calibrated for mass accuracy up to 3000 mass to charge ratio (m/z) values by using red phosphorus (MW=30.974), which can form clusters of 1-89 such molecules by laser desorption ionization in positive mode(*13*).

To analyze cholesterol distribution in the tissues by image analysis, the raw data were acquired and processed automatically by the HDI software into a collection of images. Each image corresponds to a specific m/z ion distribution over the entire tissue. The software matches each m/z value with a particular molecule of interest by searching the human metabolome database. The preferred ion state (most abundant and stable form) of cholesterol (*1*), that is, 369.35, was assigned as the m/z value(*14*). To validate this m/z value, a normal murine brain slab was incubated for 1 hour *ex vivo* with or without 10 mM methyl-ß-cyclyodextrin (mBCD) to deplete membrane cholesterol and subjected to sample preparation and image acquisition as above (Supplementary Figure S2). Three to six biological replicates (different mice) were imaged for each treatment group. Spatial distribution of cholesterol across the brain parenchyma was observed visually from mass spectrometry imaging (MSI) of the brain slides containing metastatic lesions. Cholesterol intensity distribution was not normalized to any standard molecule, and the absolute value of each pixel was reported. The dynamic range of cholesterol was kept constant (0–50000) in all figures for direct visual comparison.

### Gene Set Enrichment Analysis

Data previously generated(*6*) were assessed for cholesterol related pathways using the gene set enrichment analysis (GSEA) method developed by Subramanian et al. (*15*) to conduct the pathway analysis (Supplemental Table 1). The pathway gene sets (C2: Canonical Pathways) were obtained from the Molecular Signatures Database version 4.0 (MSigDB version 4.0, http://www.broadinstitute.org/gsea/msigdb/index.jsp) and include genes from online pathway databases, biomedical literature, and knowledge from domain experts. GSEA was conducted using the javaGSEA Desktop Application available from http://www.broadinstitute.org/gsea/downloads.jsp#msigdb. The following parameters were used:

t-test for a metric to rank genes in descending order; gene set for a permutation type as number of samples for any phenotype was less than 7 in our analysis; and 1000 for a number of permutations. All other parameters defaults in the javaGSEA Desktop Application. Nominal p-value <5% and FDR <5% were employed as the statistical cutoff for identifying enriched pathways.

### Statistical analysis

Statistical calculations were performed in GraphPad Prism 7 and SPSS v24. All tests were two-sided, and *P* values less than 0.05 were considered statistically significant. Fisher’s exact and Cochran Armitage trend tests were used to compare rates of metastatic colonization between HDL- and vLDL-pretreated groups. The non-parametric Kruskal-Wallis test and Wilcoxon rank sum test were applied to compare tumor burden between groups. The number of brain metastatic lesions and tumor burden were compared pairwise by using the non-parametric Dunn’s multiple comparisons test. In the combination WBR + simvastatin experiments, inter-rater reliability for tumor burden and number of brain metastatic lesions data were calculated by using the two-way random intraclass correlation coefficient (ICC) for average measures. Binary incidence of brain metastasis was compared by using the Fisher-Freeman-Halton test separately for each rater’s observation. Inter-rater reliability for brain metastasis incidence was calculated with the Fleiss Kappa coefficient for binary categorical measures.

## RESULTS

Analyzing previously published gene expression data from brain seeking sublines of MDA-IBC3 cells(*6*) we find pathways for cholesterol biosynthesis were three of the top five overexpressed canonical pathways in the brain *versus* lung MDA-IBC3 sublines (Supplemental Table 1), and six enzymes involved in cholesterol biosynthesis were >3-fold higher in brain than in lung sublines. This suggests that IBC3 cells in brain are depleted of cholesterol and may be upregulating cholesterol biosynthesis for survival. This supposition is consistent with recent results from Jin *et al.* on the metastatic potential of more than 500 bar-coded cell lines after intracardiac injection, which showed that brain metastatic cell lines have a lipid biosynthesis profile and require sterol regulatory element binding transcription factor 1, a transcription factor required for lipid homeostasis to survive in brain tissue(*16*). This led us to examine the effect of lipoprotein and statin treatment on IBC brain colonization and radiation response.

### Pretreating IBC3 cells with vLDL resulted in larger brain metastases at 8 weeks after injection

In a study of statin use in IBC patients, evidence of prognostic significance of statin use after post-mastectomy radiation was lost if the lipoprotein levels were known and included in the model(*8, 9*). In those studies, HDL was independently prognostic for improved outcomes after radiation; in preclinical studies, HDL and vLDL had opposite effects on radiation sensitivity(*9*). Here, to examine the effects of lipoprotein on the metastatic potential of IBC3 cells, we used a previously established tail vein–injection breast cancer metastasis model(*6*). We pretreated MDA-IBC3 cells for 24 hours *in vitro* with serum-containing or serum-free, vLDL-supplemented or HDL-supplemented, media before tail-vein injection; at 8 weeks after injection, the mice were killed and evaluated for brain metastases (Fig. 1). Percentages of mice that developed metastases in each group were 40% for control with serum, 60% for control without serum, 65% for vLDL, and 61% for HDL groups. We saw no significant differences in the number of brain metastases per mouse by treatment group (Fig. 1A). However, when we evaluated tumor burden (area positive for GFP) between groups, the vLDL group had significantly larger brain tumor burden than the HDL group (Fig. 1B, *P*=0.03, Wilcoxon rank sum test). This suggests that extra-cranial exposure to lipoproteins can alter MDA-IBC3 cells to affect proliferation of these cells as colonies in the brain.

**Figure 1.**
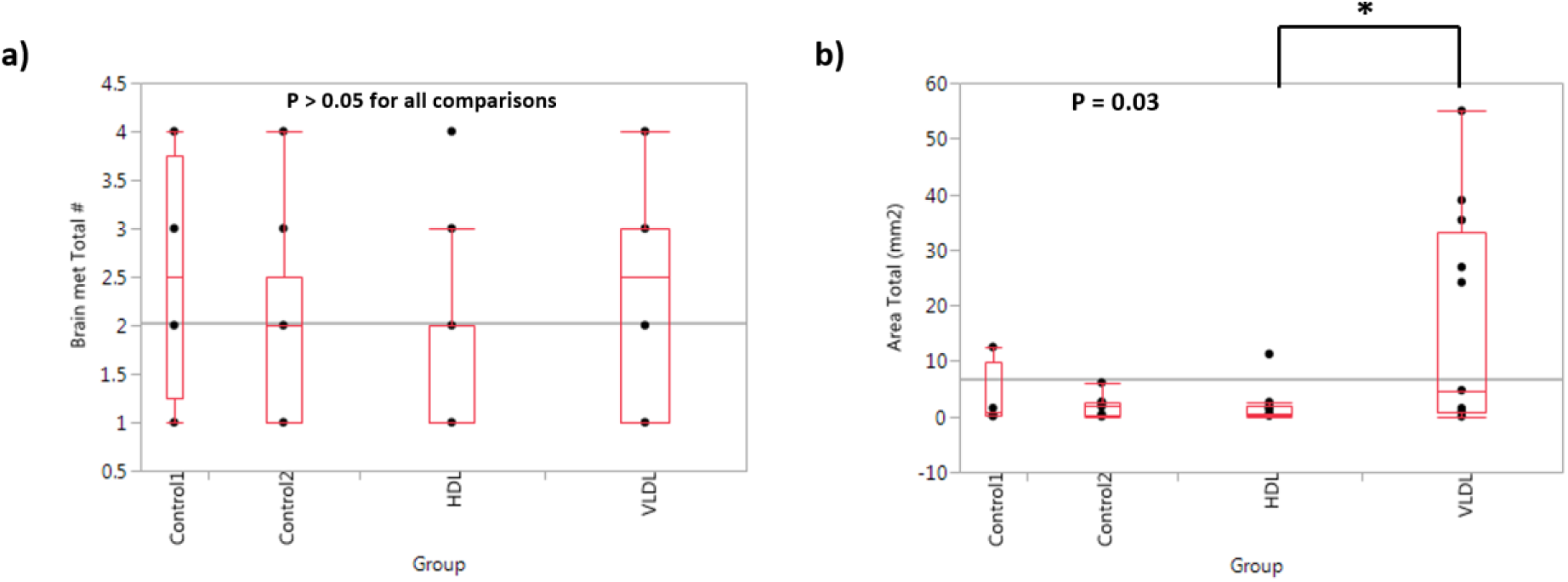
Ex Vivo treatment of MDA-IBC3 cells with vLDL increased in vivo brain metastasis burden. MDA-IBC3 cells were incubated for 24 hours in standard (control 1) or serum free media (control 2), high density lipoprotein (HDL), or very low-density lipoprotein (vLDL). 500,000 cells form each group were tail vein injected into scid/beige mice to initiate brain metastases. (A) Incidence of brain metastases per animal (yes or no). (B) Tumor burden, a global measure of GFP signal from each brain.

### Statin treatment stalled the G1 to S transition and led to impaired DNA double-strand break repair by homologous recombination and non-homologous end joining

To measure the effect of cholesterol regulation on treatment on HR and NHEJ in IBC cells, we stably integrated the TLR reporter into KPL4 cells (metastasizing to lung) and MDA-IBC3 cells (metastasizing to brain) by lentiviral transduction and used the TLR assay(*12*) to directly measure the frequency of HR and NHEJ at genomically targeted, endonuclease-induced DNA DSBs. It has been demonstrated that radiosensitization by statin is correlated with changes in the efficiency of DNA repair(*9, 11, 17*). Specifically, the observation that simvastatin treatment led to persistent H2AX ionizing radiation–induced foci^9^ led us to hypothesize that simvastatin impairs DNA double-strand break (DSB) repair in IBC cells. In mammals, the DSB repair process takes place through two major pathways, HR and NHEJ. To measure the effect of simvastatin treatment on HR and NHEJ in IBC cells, we stably integrated the TLR reporter into KPL4 cells (metastasizing to lung) and IBC3 cells (metastasizing to brain) by lentiviral transduction and used the TLR assay(*12*) to directly measure the frequency of HR and NHEJ at genomically targeted, endonuclease-induced DNA DSBs. We were able to validate detection of both HR and NHEJ repair events in both the KPL4^TLR-BFP^ and IBC3^TLR-BFP^ cell lines (Fig. 2A).

**Figure 2.**
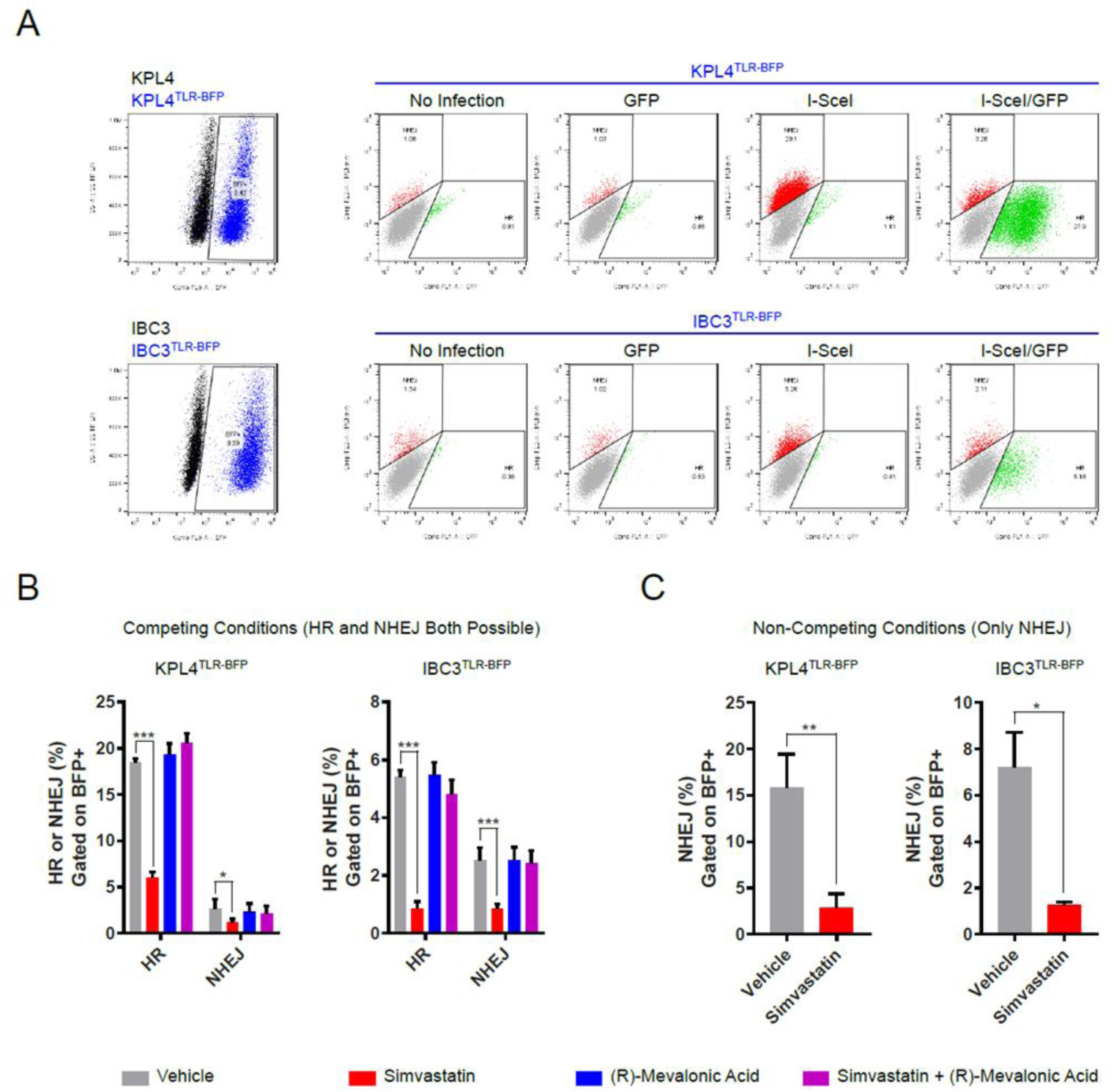
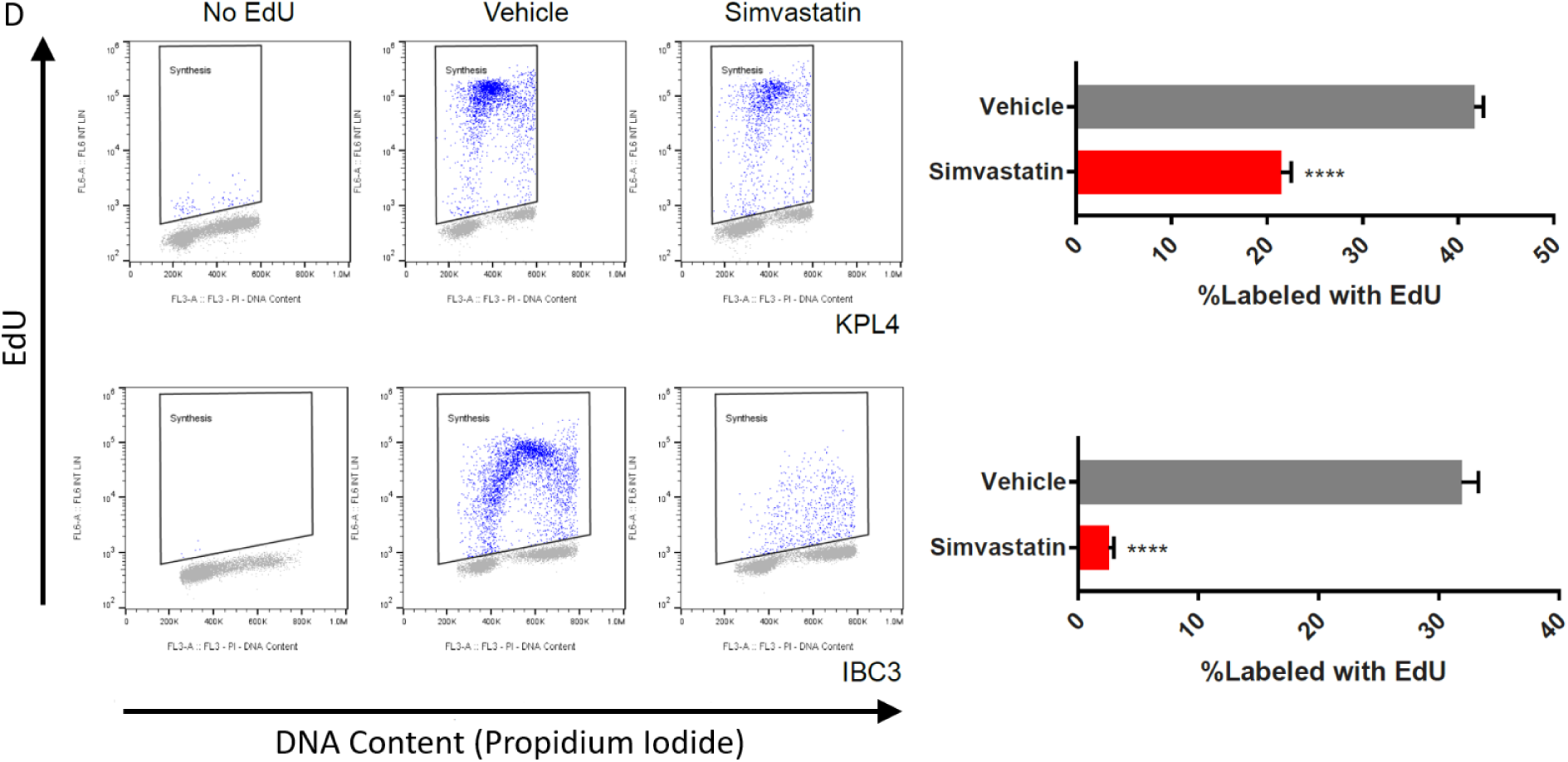
Statin treatment impairs homologous recombination (HR) and non-homologous end-joining (NHEJ) in IBC cells and suppresses cell cycle. (A) Controls showing performance of the Traffic Light Reporter (TLR) in KPB4 and IBC3 cells. (B) Frequency of HR and NHEJ events in KPB4TLR-BFP and IBC3TLR-BFP cells pre-treated with vehicle, simvastatin, (R)-mevalonic acid, or simvastatin plus (R)-mevalonic acid under competing conditions. (C) Frequency NHEJ events in KPB4TLR-BFP and IBC3TLR-BFP cells pre-treated with vehicle or simvastatin in non-competing conditions. *P<0.05, **P<0.01, ***P<0.001. (D) Pre-treating synchronized (serum-starved) KPB4 and IBC3 cells with simvastatin prior to release into serum suppresses EdU incorporation. ****P<0.001.

Pretreatment with simvastatin significantly reduced the frequency of both HR and NHEJ events when the assay was done under competing conditions (i.e., by using a lentivirus that expresses the I-SceI endonuclease and provides the homologous GFP template, thus allowing both HR and NHEJ). Importantly, treating simvastatin-treated cells with (R)-mevalonic acid (the biosynthetic intermediate produced by HMG-CoA reductase) completely rescued both HR and NHEJ (Fig. 2B). Reasoning that the competing-assay conditions may have blunted the NHEJ observations, we also performed the assay under non-competing conditions, by using a lentivirus that expresses only the I-SceI endonuclease, without providing a homologous template for HR. We again observed that simvastatin treatment strongly abrogated NHEJ in both KPL4^TLR-BFP^ and MDA-IBC3^TLR-BFP^ cell lines (Fig. 2C). These results suggest that mevalonic acid is required for the efficient repair of DNA DSBs by both HR and NHEJ, and that statins can inhibit both pathways of DSB repair by inhibiting HMG-CoA reductase-dependent mevalonic acid biosynthesis.

Repair of radiation-induced DNA damage and intrinsic radiosensitivity are highly dependent on cell cycle distribution and cell cycle checkpoint integrity. NHEJ can occur during any phase of the cell cycle, as this pathway involves direct re-ligation of DNA DSBs without requiring a template. Although non-allelic HR can occur during any phase of the cell cycle, allelic HR is the major homology-mediated pathway for repair of mammalian DSBs in somatic cells, and it requires the presence of a sister chromatid. Allelic HR is therefore only possible during S- and G2 phases. We previously showed that treating IBC cells with simvastatin reduces the fraction of cells in S phase.^11^ Because we also observed a significant reduction of HR frequency in IBC cells treated with simvastatin, we hypothesized that simvastatin may affect radiosensitivity and functional DNA repair by preventing the G1-to-S phase transition. To test this hypothesis, we synchronized KPL4 and MDA-IBC3 cells in G0/G1 by serum starvation, pretreated them with vehicle or simvastatin, and then released them into medium containing serum, EdU, and colcemid. Simvastatin significantly reduced the cumulative 24-hour incorporation of EdU in all four cell lines, suggesting that inhibition of HMG-CoA reductase 1 stalls the G1-to-S phase transition (Fig. 2D).

### Co-treatment with statin plus whole-brain radiation (10 Gy in 1 fraction) reduced the incidence of brain metastasis (number of mice with any detectable metastases) assessed 2 weeks after WBR

Using the HMG-CoA reductase inhibitor simvastatin to study the radiosensitivity of brain cholesterol regulation *in vivo* we performed four in vivo experiments.

Supplementary Figure 1 provides details of four experiments (WBR with or without statin). Therapeutic use of simvastatin is associated with cognitive side effects, and its penetrance in the brain has been estimated, (*18, 19*) and recently directly measured(*20*) demonstrating efficacy in cholesterol regulation in the brain. The simvastatin dose we used for our *in vivo* studies was based on prior in vivo experiments(*11*). <u>MDA-</u>IBC3 cells were injected into the tail vein of SCID/Beige mice, and 3 weeks later mice were assigned to one of four treatment groups. WBR dose (10 Gy in a single fraction; equivalent to 23 Gy in 2 Gy fractions with anα/β ratio of 4 in tumor) was delivered to half of the mice, and concurrent simvastatin treatment was begun in the combination WBR+statin group. (Dose-finding studies revealed acceptable toxicity and response for this radiation dose [data not shown]). Mice were sacrificed and assessed for brain metastases by stereofluoroscopy ten days after WBR. The brain metastasis incidence (number of animals with any metastases) was significantly different between groups (*P*=0.002, Cochran trend test) at that time (Fig. 3A). Representative stereoscopic brain images are shown in Figure 3B.

**Figure 3.**
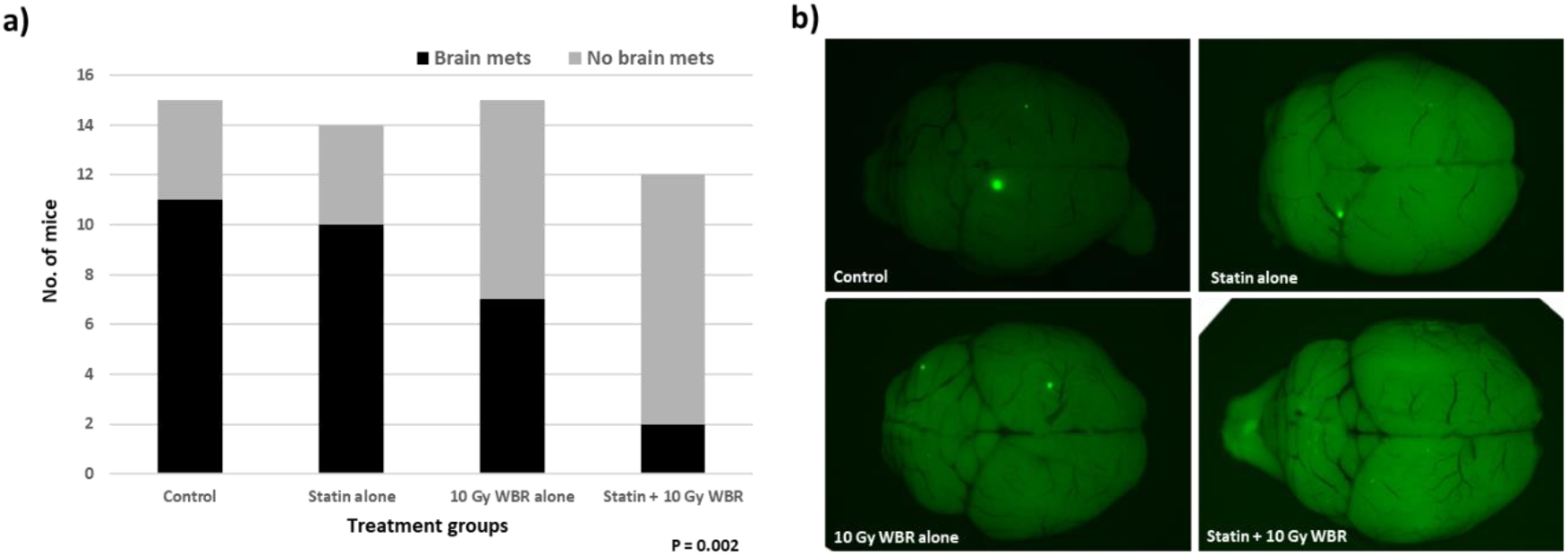
Statin + whole brain therapy decreases number of mice with brain metastases 10 days post WBR. 500,000 MDA-IBC3 cells from each group were tail vein injected into scid/beige mice to initiate brain metastases. At three weeks statin was added to the water of two group and WBR 10 Gy/1 fraction delivered. (A) Brain lesions were quantified ten days after WBR. (B) Representative stereofluorescent gross brain images.

### Co-treatment with statin plus whole-brain radiotherapy (10 Gy in 1 fraction) does not reduce brain metastasis incidence, burden, or number of metastases assessed 5 weeks after tumor initiation

Previous work on the kinetics of tail vein–injected MDA-IBC3 colonization of the brain(*21*) suggested that whole brain radiation-induced cell kill is sufficient to reduce the incidence of detectable brain metastases when the radiation is administered 5 days after injection, that is, targeting individual or small lesions, but not when macrometastases have become established (at 4 weeks after initiation). Because kinetics modeling predicted an exponential growth phase between 3 and 4 weeks(*22*), we expected a rapid transition to macrometastases at 4 weeks. To further examine the effects of WBR plus statin therapy, we initiated radiation 4 weeks after tail-vein injection, and planned evaluation at 5 weeks after irradiation (versus 2 weeks as described above) to determine the durability of this early response (Fig. 4).

**Figure 4.**
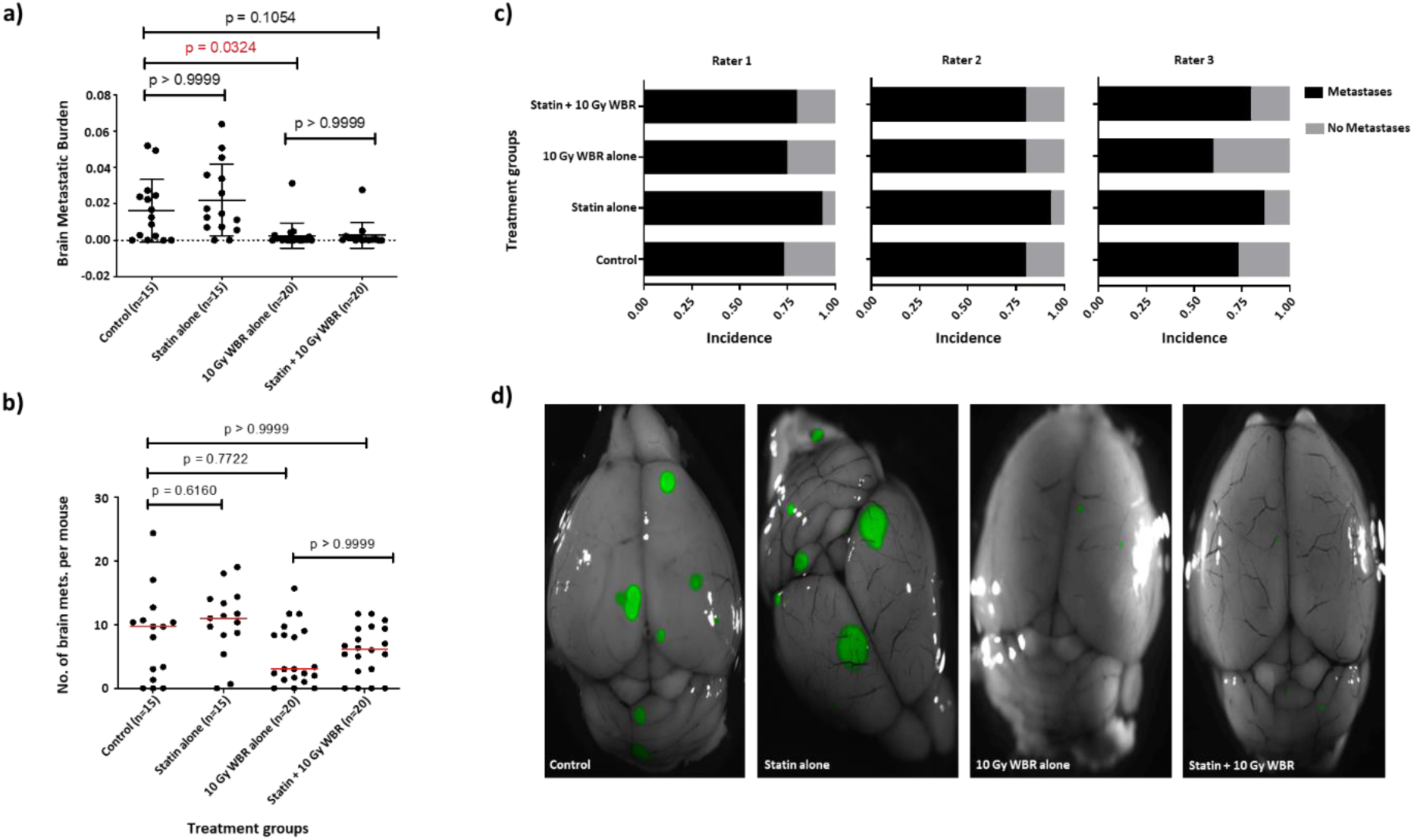
Statin + whole brain therapy (10 Gy in 1 fraction) does not reduce number of lesions or burden of brain mets compared to radiation alone five weeks after WBR. (A) Tumor burden for the no-treatment, simvastatin-alone, 10 Gy WBI–alone, and simvastatin–plus–10 Gy WBI treatment groups. Dunn’s multiple comparisons test was used to calculate the p-value for each pair of treatment groups. (B) Number of brain metastatic lesions for no-treatment, simvastatin-alone, 10 Gy WBI–alone and simvastatin–plus–10 Gy WBI treatment groups. Each dot represents a mouse. Red lines represent the median number of brain metastatic lesions in each treatment group. Dunn’s multiple comparisons test was used to calculate the p-value for each pair of treatment groups. (C) Brain metastasis incidence for no-treatment, simvastatin-alone, 10 Gy WBI–alone and simvastatin–plus–10 Gy WBI treatment groups. The Fisher-Freeman-Halton test was used to calculate the p-value for each rater (Rater 1 : p = 0.49, Rater 2 : p = 0.69, Rater 3 : p = 0.30) separately. “Fleiss’ Kappa coefficient for binary categorical agreement between three raters is 0.77, which indicates substantial agreement. (D) Representative fluorescence–bright field overlay images of no-treatment, simvastatin-alone, 10 Gy WBI–alone and simvastatin–plus–10 Gy WBI treatment groups. Each of the four images corresponds to the mouse with the tumor burden and number of brain metastatic lesions closest to the arithmetic mean value from its respective treatment group. The bright field image was overlaid with the GFP image to get both the architecture of the brain and its metastatic lesions.

Weight loss and dermatitis were observed in all 40 irradiated mice (20 each in radiation-only and radiation+statin group.) Dermatitis in the radiation field was treated with an ophthalmic ointment. Weight loss was managed with gel-pack nutrient treatment, and mice gained weight slightly after the treatment. At five weeks after WBR, simvastatin alone did not reduce any of the three endpoints: total burden of brain disease (Fig. 4A), number of brain metastases (Fig. 4B), or number of animals with brain metastases (Fig. 4C, all *P* >0.05). Representative images are shown in Figure 4D.

WBR alone (10 Gy) significantly reduced the brain metastasis burden compared with no treatment (Fig. 4A, P=0.0324) but did not reduce the number of metastases or number of mice with metastases (both *P* >0.05, Fig. 4B, C). Because WBR alone was capable of reducing tumor burden, comparisons between WBR and statin + WBR were underpowered for this endpoint (Fig. 2A). For the other endpoints (incidence and number of lesions), the effects of simvastatin and 10 Gy WBR were no different than the control condition (*P*=0.1054, Fig. 4B,C). With regard to lesion scoring from the three independent reviewers, the inter-rater reliability for number of metastases, measured with the Fleiss Kappa coefficient for binary categorical measures, was 0.77, indicating substantial agreement between the three raters. Agreement between the three raters for average measures such as the number of brain metastatic lesions, calculated with two-way ICC, was found to be 0.925 with a 95% confidence interval of 0.869– 0.956. An ICC >0.9 is considered excellent agreement between raters.

### Co-treatment with statin and fractionated whole-brain radiotherapy (3 Gy in 3 fractions) does not reduce brain metastasis incidence, burden, or number of metastases at 5 weeks

Given these results, we repeated the above *in vivo* experiment to determine if fractionated radiation, that is, a total dose of 9 Gy given in three 3-Gy fractions (equivalent to 10.5 Gy in 2-Gy fractions with an α/β ratio of 4 in tumor) would act synergistically with statins, provide a greater range to assess metastatic burden, and better mirror the clinical effects of WBR. We began statin treatment 1 week before radiation to ensure adequate loading before WBR. Still, neither statin alone nor WBR nor combination therapy reduced the burden, number, or incidence of metastases (Fig. 5). Again, inter-rater reliability, measured with the Fleiss Kappa coefficient for binary categorical measures, was 0.824 (95% CI 0.690–0.957), indicating near-perfect agreement between the three raters.

**Figure 5.**
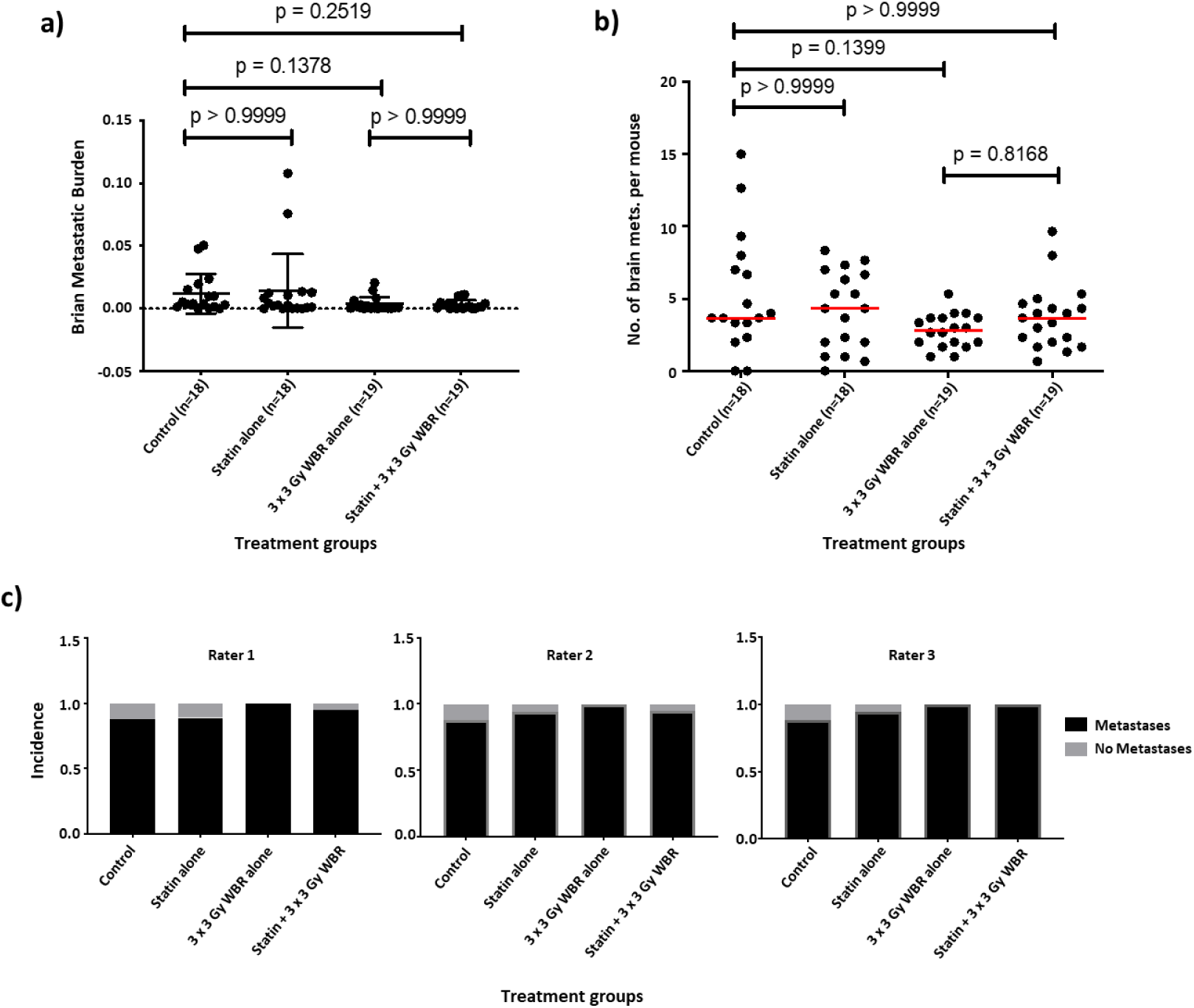
Statin + WBR (3 Gy x 3 fractions) does not reduce number of lesions or burden of brain metastases compared to radiation alone 5 weeks post WBR. (A) Tumor burden for no-treatment, simvastatin alone, 9 Gy total WBI dose given in 3 fractions alone, and the combination group. Dunn’s multiple comparisons test was used to calculate the p-value for each pair of treatment groups. (B) Number of brain metastatic lesions for groups of no-treatment, simvastatin alone, 9 Gy total WBI dose given in 3 fractions alone, and combination treatment. Each dot represents a mouse. Red lines represent the median number of brain metastatic lesions in each treatment group. Dunn’s multiple comparisons test was used to calculate the p-value for each pair of treatment groups. (C) Brain metastasis incidence for no-treatment, simvastatin alone, 9 Gy total WBI dose given in 3 fractions alone, and the combination group. Fisher-Freeman-Halton test was used to calculate the p-value for each rater ((Rater 1: p = 0.47, Rater 2: p = 0.51, Rater 3 : p = 0.25) separately. Fleiss’ Kappa coefficient for binary categorical agreement (yes vs. no) is 0.824 (95% CI 0.690-0.957).

### Mass spectrometry imaging of postmortem brain tissues reveals global increase in brain cholesterol after irradiation

Although we sought to evaluate the simvastatin target HMG CoA reductase and cholesterol synthesis in the in vivo metastases, we found that that was not feasible because of the low tumor burden in irradiated samples. For these experiments, we generated whole-brain lysates from samples from each group (n=3 brains per group) and subjected them to HMG CoA reductase immunoblot analysis (Fig. 6A). Variability between brain samples in the same group was evident. Although this analysis is limited by the small number of samples, radiation increased HMG CoA reductase expression in the brain compared with control (*P*=0.04), but HMG CoA reductase was unchanged after simvastatin + WBR versus control (Fig. 6B). To directly assess cholesterol in the normal brain and tumor lesions, we used mass spectrometry imaging (MSI) of whole-brain tissue sections adjacent to H&E-stained sections to enrich for sections containing metastases. To validate the published m/z ratio for cholesterol, a slice of normal brain was incubated *ex vivo* with mBCD to chelate cholesterol from the tissue. MSI analysis revealed that significant cholesterol was present in the *ex vivo* control section and was substantially reduced by mBCD (Fig. 6C). In both untreated and statin-alone-treated brains, the signal intensity for cholesterol was below the level of detection in brain and, where evaluable, in metastases (Fig. 6C). The intensity heat scale was held constant across groups for comparison. In the radiation-alone and radiation + statin brains, the cholesterol signal was evident in normal brain, suggesting increased synthesis after radiation consistent with the increased expression of HMG Co A reductase noted above. Signal in individual lesions, however, remained below the threshold of detection and thus remained inadequate to demonstrate or refute the effectiveness of statin on the tumor itself. Although the analysis was underpowered, tumor burden did not correlate with cholesterol levels in the brain parenchyma or its metastatic regions in any of the four treatment groups (not shown).

**Figure 6.**
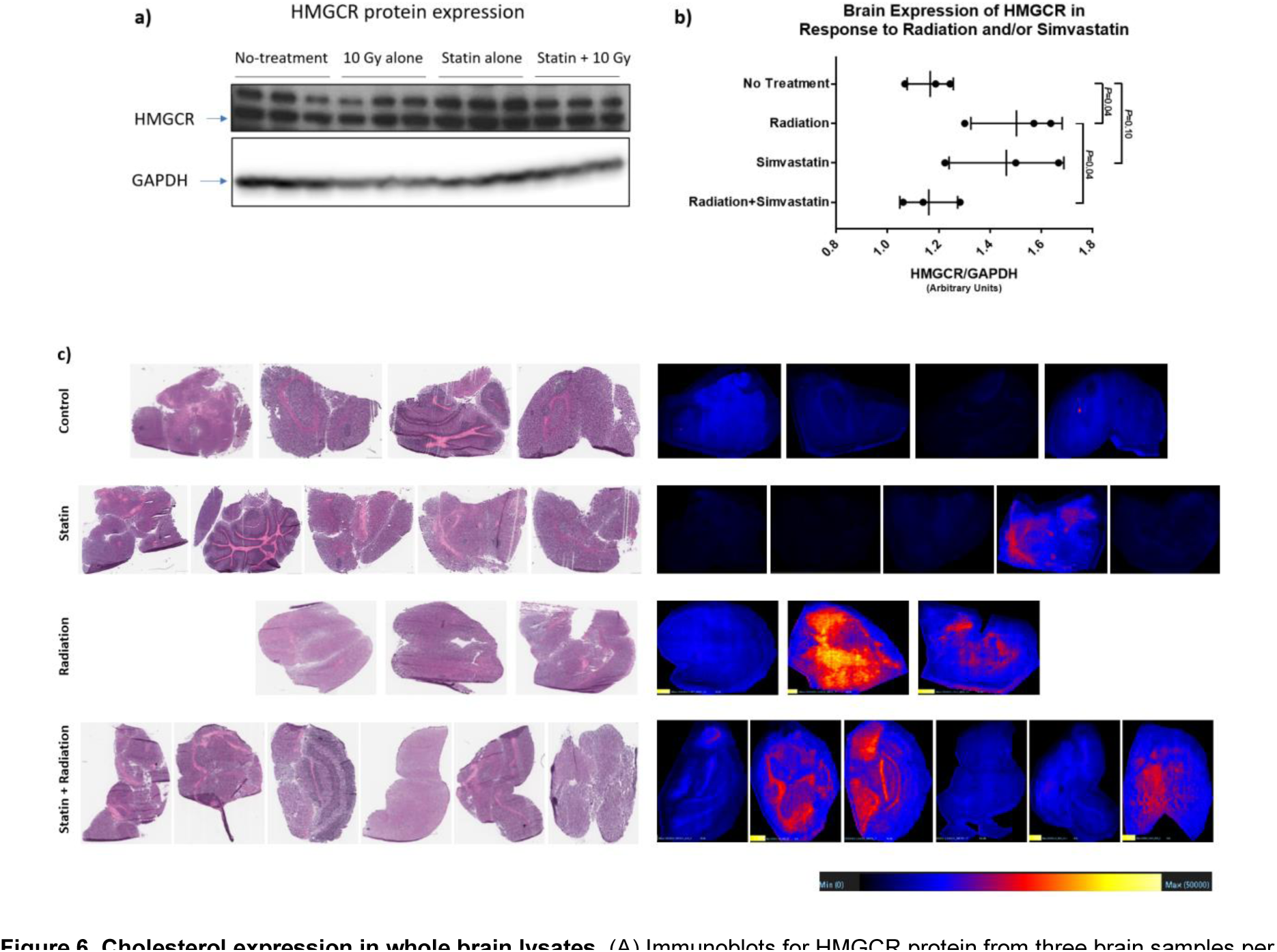
Cholesterol expression in whole brain lysates. (A) Immunoblots for HMGCR protein from three brain samples per group. (B) Densitometry quantification of A. (C) Cholesterol distribution in the brain parenchyma using mass spectroscopy imaging. Scale bar of intensity heat maps were kept constant across all four treatment groups (0–50000) to visually compare the cholesterol distribution. Serial hematoxylin and Eosin (H and E) and MSI images m/z value, 369.35.

## Discussion

Several lines of evidence, including RNA expression data from MDA-IBC3 brain sublines (Supplemental Table 1), suggest that cholesterol regulation has a role in the survival of brain tumors and brain metastases(*16, 23, 24*). Clinical and preclinical studies point to synergy between cholesterol regulation and radiation, particularly in aggressive breast cancer subtypes(*8, 9, 11, 17*). Here we report that in a HER2^+^ IBC model of brain metastases, tumor burden was enhanced after *ex vivo* incubation with cholesterol-loading vLDL, and we describe early signs of a signal for radiation and statin synergy in small lesions with short follow-up.

However, we found no evidence of durable synergy from treating MDA-IBC3 macrometastases, and found an unexpected increase in HMG CoA Reductase expression and cholesterol in normal brain tissues in response to radiation without evidence of suppression of HMG CoA Reductase in brain metastases from animals in statin groups. Our results suggest that in this model radiation effectiveness is greater when used to treat smaller lesions, which we have also shown in previous modeling studies(*22*). The extracranial experiments support the idea that extracranial lipoproteins may influence the survival of cells in the brain. Still, limitations notwithstanding we find no evidence of durable synergy between statin and WBR for established brain metastases.

Development of brain metastases involves several biological steps for cells to escape the primary site, survive in the circulation, escape into the metastatic site, and grow and survive in the metastatic environment. Our tail-vein injection model for establishing brain metastases captures survival in the circulation, passage through the lungs, and colonization in the brain, but not escape of cells from the primary tumor site(*25*). Cardiac-injection models can capture similar steps but may exclude passage through the cardiopulmonary system to reach the brain. Jin *et al* reported cardiac injection of more than 500 barcoded cell lines and RNA sequencing of brain metastases in 7 of 21 breast cancer cell lines that develop metastases(*16*); 4 of those lines contained phosphatidylinositol-3-kinase (PI3K) mutations that are downstream of HER2. Gene signatures that were increased in brain metastases implicate signaling from PI3K, HER2, and adipogenesis, but signatures associated with fatty acid metabolism were decreased. Knocking out SREBF1, the top differentially expressed gene in the brain, significantly reduced brain lesions. Our work comparing brain-metastasizing to lung-metastasizing sublines from HER2^+^ MDA-IBC3 cells is consistent with these reports highlighting lipid metabolism as a top expression pathway and emphasize the importance of testing cholesterol-targeting strategies to treat brain metastases.

Statins are prescribed to millions of individuals worldwide and have a well-established safety profile. Among the lipophilic statins expected to cross the BBB, simvastatin, pitastatin, and lovastatin are also radiosensitizers(*17*) that have similar effects on breast cancer cell lines *in vitro*, whereas atorvastatin, pravastatin, and rosuvastatin are not(*17*). A direct assessment of simvastatin in the brain estimated >25% BBB penetration of simvastatin versus <5% of atorvastatin(*26*), and clinical studies have shown changes in biomarkers in the cerebrospinal fluid from oral statins at clinically used doses(*27*). Most recently, plasma and brain levels of statins as well as lipid regulation in primary brain cancer models demonstrate brain penetration by statins. Together, these studies highlight the value of studying oral simvastatin and synergy between simvastatin and radiation in the current study.

Herein, we observed enhanced progression of brain lesions from cells that had been incubated with vLDL *ex vivo*, which suggests that extracranial lipoprotein levels may participate in loading circulating cells that will survive in the brain or alter signaling that drives brain proliferation. Although our findings from early lesions (those present at 2 weeks after irradiation) suggest that simvastatin plus radiation can reduce the incidence of those early lesions, examination at 5 weeks after irradiation demonstrates no effect on incidence of lesions, number of lesions, or lesion burden in this model. As predicted in previous mathematical models of brain metastases and effect of radiation from this cell line (*22*), a minimally adequate dose of radiation (10 Gy in 1 fraction) can reduce the size of each metastases (assessed as burden) but does not cure mice; the incidence of brain metastases–bearing mice is not reduced. Indeed, we previously showed that the effectiveness of radiation in the brain is directly related to intrinsic radiation sensitivity of cells and was curative only against microscopic disease (*21*). In the current study, the lower biologically effective dose of 9 Gy in three 3-Gy fractions did not even reduce burden. However, we used these subclinical doses to assess possible synergy with simvastatin. Indeed, we found that irradiation 3 weeks after tumor initiation, before exponential growth (as suggested by modeling studies) demonstrated an early signal for reduced tumor burden 2 weeks later, suggesting enhanced efficacy of radiation at that time; however, irradiation just after this growth curve at 4 weeks, assessed for durability 5 weeks later, showed no difference in benefit, whether from single-dose radiation or fractionated radiation. The implications of an increase in total brain cholesterol after radiotherapy may warrant further investigation for its effects on normal tissue toxicity.

We acknowledge several limitations of the current study. First, we used a tail-vein-injection model rather than a spontaneous breast cancer model, meaning that a single bolus of breast cancer cells entered the circulation rather than being shed from the primary or metastatic sites over time; moreover, only single cells entered the circulation, which may not fully recapitulate the clinical situation(*28–30*). Nevertheless, in this model, the effect of statins on cells established in the brain (rather than extracranial cells before they enter the brain) can be largely isolated. Second, we examined this hypothesis in only one animal model. However, the similarities between our expression data (Supplemental Table 1) and the cell lines analyzed by Jin *et al*. are encouraging their possible generalizability(*16*). An additional limitation is the immunocompromised state inherent in all xenograft models. Efimova *et al.* noted synergy between radiation and pitastatin *in vivo*, but that was in an immunocompetent melanoma model(*17*). No syngeneic mouse models derived from IBC cells exist at this time. Finally, without direct analysis of brain concentrations of simvastatin, we cannot confirm penetrance of simvastatin into the brain metastases, however we note these studies have been performed by others and suggest simvastatin can penetrate the brain(*6*), although direct penetrance into brain metastases is not demonstrated here.

In conclusion, we report gene expression data suggesting that cholesterol signaling is increased in brain metastases derived from HER2^+^ MDA-IBC3 cells injected in the tail vein as compared with lung metastases. Although the current study showed that *ex vivo* incubation with vLDL increased the burden (growth) of brain lesions and that early concurrent statin plus WBR reduced the burden of brain disease observable at 2 weeks after single-dose radiation, an experiment treating lesions grown for a longer period and observed at longer intervals after treatment (which are more compatible with clinical translation) showed no evidence of durable benefit with regard to incidence or tumor burden. The purported radiation-sensitizing signal observed after 2 weeks is interesting but presumed to be transient at best using this approach. Subsequent studies with agents having better BBB penetrance, such as liposomes(*31*) or smaller, more bioavailable targeted agents, may be preferable.

## Acknowledgements

We are thankful to Dr. Swathi Arur for use of the fluorescent stereomicroscope. We would like to thank Charlie Kingsley for assistance in operating the XRAD, Jie Zhang for sectioning assistance, Dodge Lo Baluya for MSI assistance and Dr. Amy Ninetto (of MD Anderson’s Research Medical Library) and Christine Wogan (of MD Anderson’s Division of Radiation Oncology) for editing the manuscript.

