## Supplemental Material for "*In Vivo* Simvastatin and Brain Radiation in a Model of HER2^+^ Inflammatory Breast Cancer Brain Metastasis"

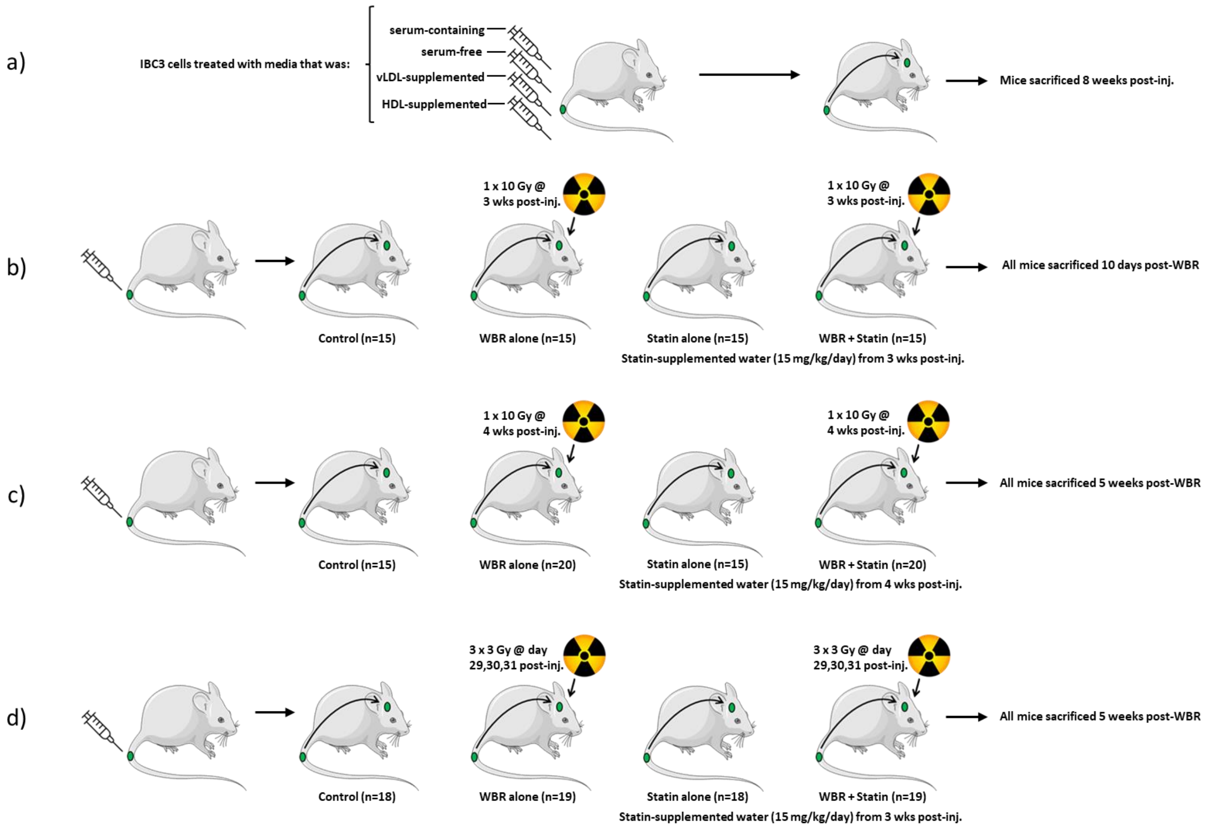


**Supplementary Figure S1. Experimental schema.** GFP-labeled MDA-IBC3 cells (500,000 per mouse) were injected intravenously through the tail vein of SCID/Beige mice in all cases. (A) For Experiment 1, IBC3 cells were incubated *in vitro* as described and injected by tail vein. Mice were killed (by $CO_2$ inhalation and vertebral dislocation) for evaluation at 8 weeks after injection. (B) For Experiment 2, 3 weeks after tail vein injection, therapy (statin, radiation alone, or combination) was begun, and mice were killed 2 weeks later. (C) For Experiment 3, a single 10-Gy whole-brain radiotherapy dose was given at 4weeks after cell injection to 40 mice (2 groups). Statin treatment was started immediately after radiation for 35 mice (2 groups) and continued for 5 weeks. (D) For Experiment 4, three 9-Gy fractions were given at 4 weeks after cell injection to 38 mice (2 groups). Statin treatment was started for 37 mice (2 groups) at 1 week before the first radiation dose (from Day 22) and was continued for 5 weeks after irradiation. At 5 weeks after radiation treatment, mice from all four groups (in both experiments) were killed and their brains and lungs were collected to identify metastatic lesions.

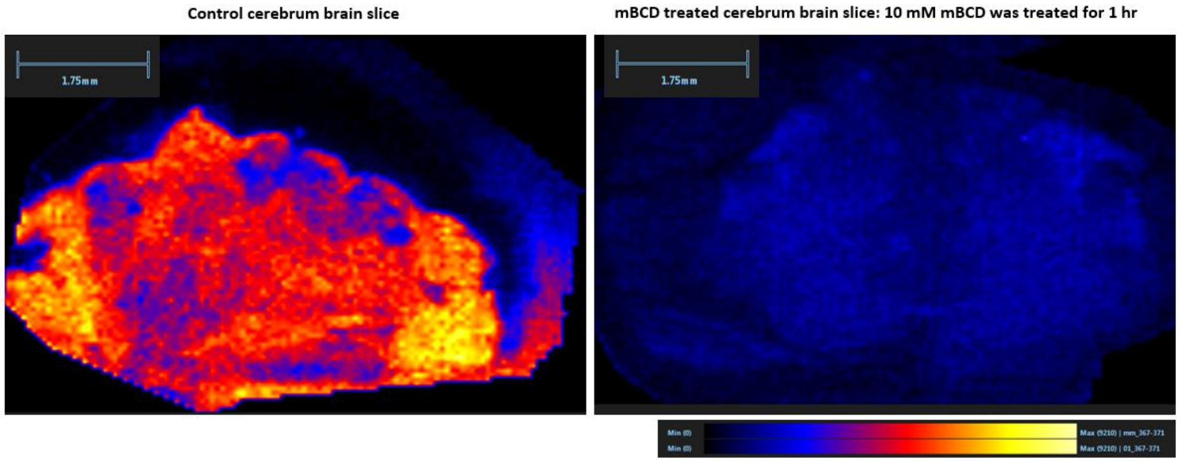


**Supplementary Figure S2. Validation of cholesterol depletion with methyl-ß-cyclyodextrin (mBCD) in mouse brain slices.** (A) Normal brain slice incubated *ex vivo* in DMEM and subjected to mass spectrometry imaging (MSI) processing and analysis for the indicated m/z value (369.35). (B) Normal brain slice incubated *ex vivo* in DMEM + 10 mM methyl-ß-cyclyodextrin (mBCD) and subjected to MSI processing and analysis for the indicated m/z value (369.35).

**Supplementary Table 1. Cholesterol regulation in brain-metastasizing versus lung-metastasizing sublines of MDA-IBC3 inflammatory breast cancer cells.** We used gene array analysis to reveal that the cholesterol synthesis pathway is differentially regulated between lung metastasis cell lines and brain metastasis cell lines (top). The main differentially regulated genes are shown at the bottom of the table.

| Top Canonical Pathways – Brain vs. Lung Sublines | | |
|---|---|---|
| **Name** | **P-Value** | **Ratio** |
| **Superpathway of Cholesterol Biosynthesis** | **$8.57 \times 10^{-7}$** | **13/27** |
| Hepatic Fibrosis/Hepatic Stellate Cell Activation | $1.69 \times 10^{-6}$ | 43/195 |
| Role of Pattern Recognition Receptors/Bacteria and Viruses | $9.08 \times 10^{-6}$ | 29/118 |
| **Cholesterol Biosynthesis** | **$1.14 \times 10^{-5}$** | **8/13** |
| **Cholesterol Biosynthesis II (via 24, 25-dihydrolanosterol)** | **$1.14 \times 10^{-5}$** | **8/13** |

| Symbol | Entrez Gene Name | P-Value | Expr Fold Change (Lung/Brain) |
|---|---|---|---|
| DHCR24 | 24-dehydrocholesterol reductase | $2.07 \times 10^{-5}$ | -3.00 |
| FDFT1 | famesyl-diphosphate famesyltransferase 1 | $2.95 \times 10^{-5}$ | -5.63 |
| HMGCR | 3-hydroxy-3-methylglutaryl-CoA reductase | $3.72 \times 10^{-7}$ | -3.85 |
| HMGCS1 | 3-hydroxy-3-methylglutaryl-CoA synthase 1 | $7.62 \times 10^{-8}$ | -8.23 |
| IDI1 | isopentenyl-diphosphate delta isomerase 1 | $6.39 \times 10^{-5}$ | -3.00 |
| MSMO1 | methylsterol monooxygenase 1 | $1.40 \times 10^{-6}$ | -3.74 |
| SC5D | sterol-C5-desaturase | $1.69 \times 10^{-6}$ | -7.91 |
| SQLE | Squalene epoxidase | $2.42 \times 10^{-7}$ | -4.02 |
